# DHTKD1 and OGDH display in vivo substrate overlap and form a hybrid ketoacid dehydrogenase complex

**DOI:** 10.1101/645689

**Authors:** João Leandro, Tetyana Dodatko, Jan Aten, Ronald C. Hendrickson, Roberto Sanchez, Chunli Yu, Robert J. DeVita, Sander M. Houten

## Abstract

Glutaric aciduria type 1 (GA1) is an inborn error of lysine degradation characterized by a specific encephalopathy that is caused by toxic accumulation of lysine degradation intermediates. Substrate reduction through inhibition of DHTKD1, an enzyme upstream of the defective glutaryl-CoA dehydrogenase, has been investigated as a potential therapy, but revealed the existence of an alternative enzymatic source of glutaryl-CoA. Here we show that loss of *DHTKD1* in GCDH-deficient HEK-293 cells leads to a 2-fold decrease in the established GA1 clinical biomarker glutarylcarnitine, and demonstrate that OGDH is responsible for this remaining glutarylcarnitine production. We furthermore show that DHTKD1 interacts with OGDH, DLST and DLD to form a hybrid α-ketoglutaric and α-ketoadipic acid dehydrogenase complex. In summary, α-ketoadipic acid is an in vivo substrate for DHTKD1, but also OGDH. The classic α-ketoglutaric dehydrogenase complex can exist as a previously undiscovered hybrid containing DHTKD1 displaying improved kinetics towards α-ketoadipic acid.

## INTRODUCTION

Lysine degradation proceeds mainly via the mitochondrial saccharopine pathway consisting of 9 different enzymatic steps that ultimately yield 2 acetyl-CoAs and several reduction equivalents (Figure S1) (Crowther et al., 2019; Pena et al., 2017; Struys and Jakobs, 2010; Struys et al., 2014). In this pathway, α-ketoadipic acid (AKA) undergoes oxidative decarboxylation catalyzed by a recently discovered α-ketoadipic acid dehydrogenase complex (KADHC). The existence of this complex is based on the identification of mutations in *DHTKD1* in individuals with α-aminoadipic and α-ketoadipic aciduria, an established inborn error of lysine metabolism (Danhauser et al., 2012; Hagen et al., 2015; Stiles et al., 2016). *DHTKD1* encodes the dehydrogenase E1 and transketolase domain-containing protein 1 (DHTKD1), a close protein homolog of OGDH, the E1 subunit of the α-ketoglutarate dehydrogenase complex (KGDHC) (Bunik and Degtyarev, 2008). The KADHC and KGDHC belong to the family of α-ketoacid dehydrogenase complexes, which consist of several E1, E2 and E3 subunits. The E1 and E2 subunits of these complexes are unique and catalyze the oxidative decarboxylation and transacylation of the substrate to form a Coenzyme A (CoA) ester. The E3 subunit is shared (dihydrolipoyl dehydrogenase (DLD)) between all complexes and regenerates the lipoamide cofactor of the E2 subunit. KADHC and KGDHC most likely share the E2 subunit (dihydrolipoyl succinyltransferase; DLST) subunit given the lack of novel candidate E2 subunits and the homology between OGDH and DHTKD1. Indeed recombinant DHTKD1, DLST and DLD are able to assemble into an active KADHC in vitro (Nemeria et al., 2018). The biochemical and enzymatic properties of the native KADHC in mammalian tissues have not been characterized.

Lysine degradation is clinically relevant pathway, because of two severe inborn errors of metabolism. Pyridoxine-dependent epilepsy is a condition characterized by severe seizures that can be treated by pyridoxine. It is caused by mutations in *ALDH7A1* (Mills et al., 2006; van Karnebeek et al., 2016). Glutaric aciduria type 1 (GA1) is a cerebral organic aciduria caused by a defect in glutaryl-CoA dehydrogenase encoded by *GCDH* (Figure S1). Patients can present with brain atrophy and macrocephaly and may develop dystonia due to striatal degeneration with acute or insidious onset (Boy et al., 2018; Goodman et al., 1998; Greenberg et al., 1995). Remarkably, hyperlysinemia and α-aminoadipic and α-ketoadipic aciduria (Dancis et al., 1983; Houten et al., 2013; Stiles et al., 2016), two other defects in lysine metabolism, are considered biochemical phenotypes of questionable clinical significance (Goodman and Duran, 2014).

Substrate reduction therapies in inborn errors aim to decrease the levels of toxic metabolite by inhibiting an enzyme upstream of the defective enzyme. The occurrence of seemingly non-harmful enzyme defects in the lysine degradation pathway suggests that substrate reduction therapy by inhibiting these enzymes is safe in humans. Accordingly, inhibition of α-aminoadipic semialdehyde synthase (AASS) has been suggested as a therapeutic intervention in pyridoxine-dependent epilepsy (Pena et al., 2017). Similarly, inhibition of DHTKD1 has been proposed as a therapeutic intervention in GA1 (Biagosch et al., 2017). This approach was tested in *Dhtkd1/Gcdh* double knockout (KO) mice. Unexpectedly, knocking out *Dhtkd1* did not appear to mitigate the clinical and biochemical phenotype of the *Gcdh* KO mouse (Biagosch et al., 2017). This result indicates that DHTKD1 is not an exclusive source of glutaryl-CoA. Here we reassessed the role of DHTKD1 in lysine degradation and identified the alternative source of glutaryl-CoA.

## RESULTS

### The Hypomorphic C57BL/6 *Dhtkd1* Allele is not Protective in a GA1 Mouse Model

The identification of *DHTKD1* mutations in individuals with α-aminoadipic and α-ketoadipic aciduria suggests that DHTKD1 is essential for the conversion of α-ketoadipic acid into glutaryl-CoA (Danhauser et al., 2012; Hagen et al., 2015; Stiles et al., 2016). Additional evidence was provided by the study of the BXD recombinant inbred mouse strains. In this population of mouse strains the *Dhtkd1*^B6^ allele associates with decreased *Dhtkd1* mRNA and DHTKD1 protein, increased plasma α-aminoadipic acid (AAA) and increased urine α-ketoadipic acid (Chick et al., 2016; Leandro et al., 2019; Wu et al., 2014). Unexpectedly, knocking out *Dhtkd1* did not appear to mitigate the clinical and biochemical phenotype of the *Gcdh* KO mouse (Biagosch et al., 2017). This experiment, however, was performed in mice on a mixed genetic background (37.5% C57BL/6, 12.5% FVB and 50% 129) (Biagosch et al., 2017). Given that both C57BL/6 and FVB are DHTKD1 deficient, we argued that a partial therapeutic effect may have been missed and decided to readdress this hypothesis taking advantage of the natural variation in the murine *Dhtkd1* locus. We crossed the C57BL/6 congenic *Gcdh* KO mouse with DBA/2J to obtain an F1 progeny. Double heterozygous (*Gcdh*^+/−^ *Dhtkd1*^D2/B6^) mice were then intercrossed to generate an experimental cohort of B6D2F2 mice. The genotype distribution in the B6D2F2 was Mendelian with no lethality of GA1 mice.

We next evaluated the consequences of the different *Gcdh* and *Dhtkd1* genotypes on plasma and urine biomarkers for GA1. The hypomorphic C57BL/6 *Dhtkd1* allele did increase plasma AAA and urine AKA, but did not decrease the accumulation of plasma glutarylcarnitine (C5DC), and urine glutaric and 3-hydroxyglutaric acid (Figure 1). This result further establishes that DHTKD1 is not the only source of glutaryl-CoA, which is consistent with the work by Biagosch et al. (Biagosch et al., 2017). Interestingly, we noted that *Gcdh* KO mice had an increase in plasma AAA and urine AKA, which appears independent of the *Dhtkd1* genotype (Figure 1). This indicates that the KADHC is partially inhibited in the *Gcdh* KO likely due to inhibition by its accumulating product glutaryl-CoA.

**Figure 1.**
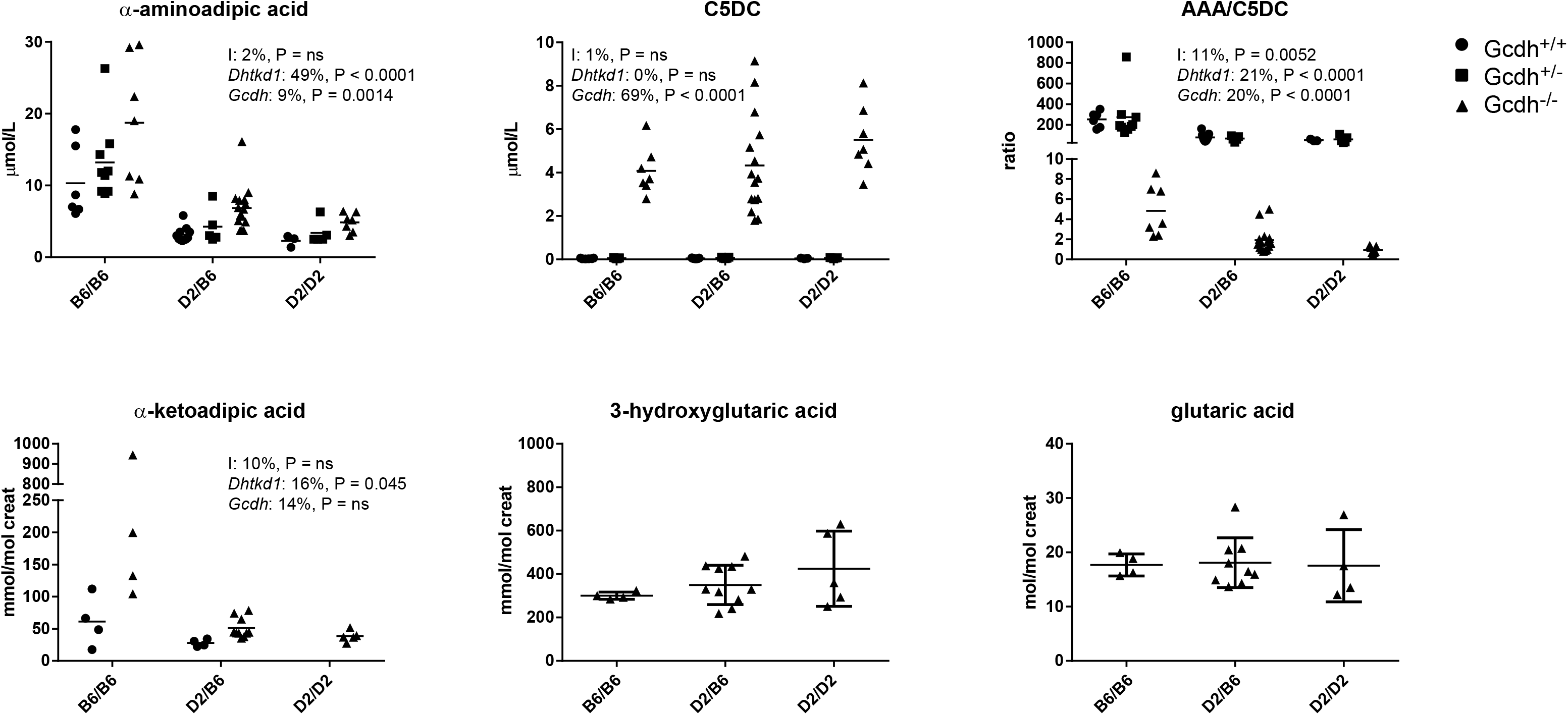
DHTKD1 Deficiency does not Limit the Accumulation of Plasma C5DC in *Gcdh* KO Mice. Biochemical analyses of plasma C5DC and AAA and urine AKA, glutaric and 3-hydroxyglutaric acids in *Gcdh* KO mice generated with wild type (DBA/2J: D2) and hypomorphic *Dhtkd1* (C57BL/6: B6) alleles. Plasma C5DC and AAA are also shown for *Gcdh*^+/+^ and *Gcdh*^+/−^ and urine AKA is also shown for *Gcdh*^+/+^. The ratio of AAA/C5DC is displayed. A two-way analysis of variance was performed and the results (% variation explained and P value) are displayed for each graph. I denotes the interaction term; *Dhtkd1* denotes the effect of the *Dhtkd1* allele (B6 or D2); *Gcdh* denotes the effect of the *Gcdh* genotype (*Gcdh*^+/+^, *Gcdh*^+/−^ and *Gcdh*^−/−^); ns, not significant. Error bars indicate SD.

### DHTKD1 Knockout Decreases Glutarylcarnitine in GCDH-deficient Cell Lines

To further investigate the enzymatic sources of glutaryl-CoA, we used CRISPR-Cas9 genome editing in HEK-293 cells. Single and double KO cell lines for *GCDH* and *DHTKD1* were generated simultaneously (Figure S2A). CRISPR-Cas9-induced mutations resulted in full KO cell lines as demonstrated by absent GCDH and DHTKD1 protein in immunoblots (Figure S2A). As predicted, single *GCDH* and *DHTKD1* KO cell lines had increased C5DC and AAA, respectively (Figures S2B and S2C). AAA was also elevated in *GCDH/DHTKD1* double KO cell lines (Figures S2C). The level of C5DC was reduced 2-fold in *GCDH/DHTKD1* double KO cell lines when compared to single *GCDH* KO cell lines. C5DC levels, however, did not decrease to control levels (Figures S2B).

The variation in C5DC concentrations was relatively large between the different *GCDH/DHTKD1* double KO cell lines, which may be related to the clonal isolation. We therefore selected 3 different *GCDH* single KO cell lines, and then targeted *DHTKD1* using CRISPR-Cas9 to generate *GCDH/DHTKD1* double KO cell lines (Figure 2A and Figure S2D). These consecutively generated *GCDH/DHTKD1* double KO cell lines reproduced the results obtained in the simultaneously generated *GCDH/DHTKD1* double KOs and showed an overall decrease in the level of C5DC. Again C5DC did not decrease to control levels (Figure 2A and Figures S2E and S2F). Of the 11 isolated *GCDH/DHTKD1* double KO cell lines 8 had decreased (between 18-60% decrease), 2 had similar and 1 had increased C5DC levels (Figure 2A). As expected the level of AAA increased in all double KO cell lines. In conclusion, these data suggest that DHTKD1 inhibition can limit metabolite accumulation in GCDH deficient cell lines, but also confirm that there is an alternative source of glutaryl-CoA that may be able to compensate partially for the loss of DHTKD1.

**Figure 2.**
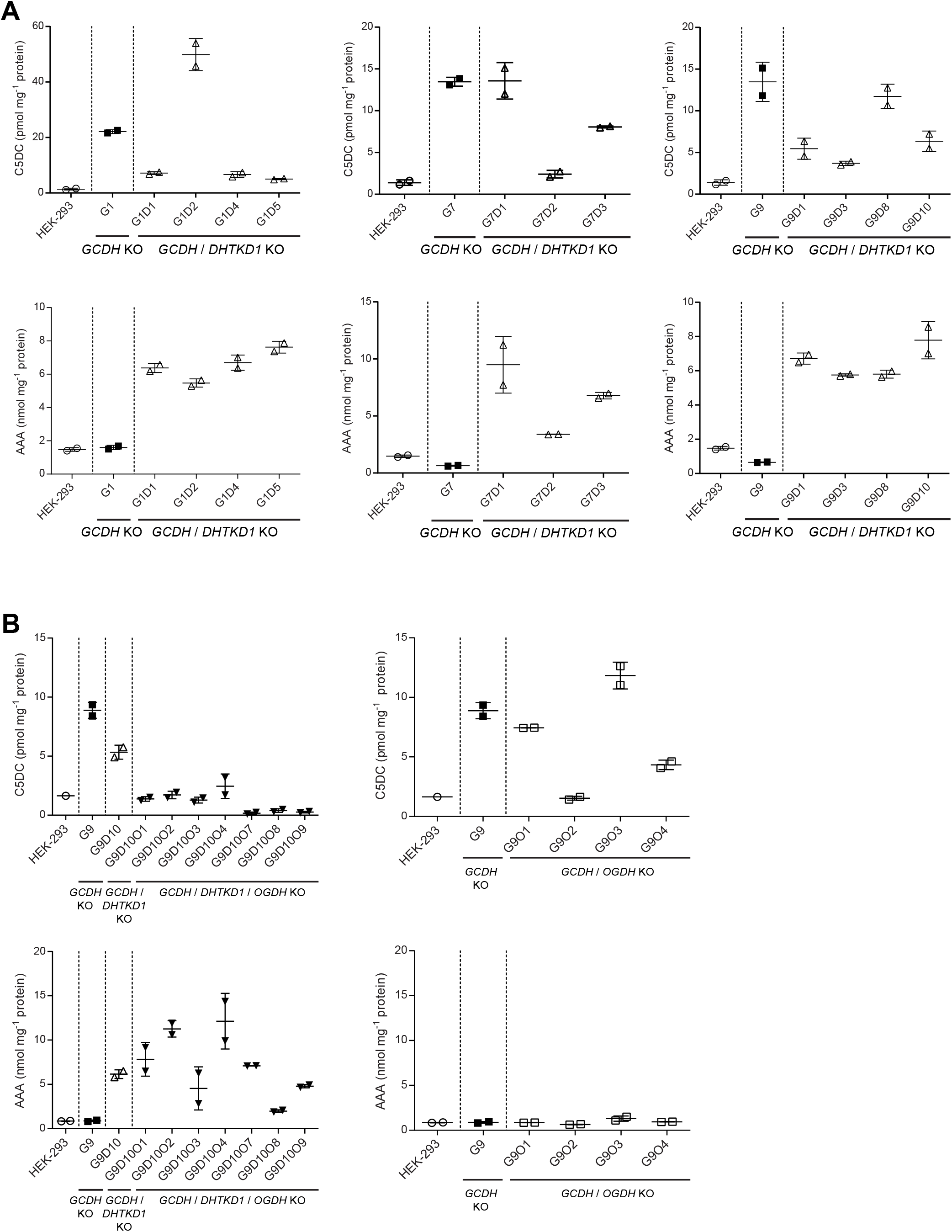
Genetic Dissection of the Source of Glutaryl-CoA in *GCDH* KO Cells. (A) Biochemical analyses of C5DC (*upper panel*) and AAA (*lower panel*) levels in HEK-293 *GCDH/DHTKD1* double KO (△) compared with HEK-293 *GCDH* single KO (■) and wild-type HEK-293 cells (◯). Single *GCDH* KO clones are indicated as G# and double *GCDH/DHTKD1* KO clones as G#D#. Three HEK-293 *GCDH* KO clones (G1, G7 and G9) were selected for the generation of the *DHTKD1* KO. Error bars indicate SD. (B) Biochemical analyses of C5DC (*upper panel*) and AAA (*lower panel*) levels in HEK-293 *GCDH/DHTKD1/OGDH* triple KO (▼) and HEK-293 *GCDH/OGDH* double KO (●) compared with HEK-293 *GCDH/DHTKD1* double KO (△), HEK-293 *GCDH* single KO (■) and wild-type HEK-293 cells (◯). Single *GCDH* KO clones are indicated as G#, double *GCDH/DHTKD1* KO clones as G#D#, double *GCDH/OGDH* KO clones as G#O# and triple *GCDH/DHTKD1/OGDH* KOclones as G#D#O#. The HEK-293 *GCDH/DHTKD1* double KO clone G9D10 and HEK-293 *GCDH* single KO G9 were selected for the generation of the *OGDH* KO. Error bars indicate SD.

### OGDH Partially Compensates for the Loss of DHTKD1

Enzyme analysis revealed that purified KGDHC from porcine heart is able to use AKA albeit with a lower affinity and at a lower rate when compared to α-ketoglutaric acid (AKG) (V_max, AKG_ = 26.33 μmol·min^−1^·mg^−1^, V_max, AKA_ = 6.7 μmol·min^−1^·mg^−1^ and *K*_m AKG_ = 249 μM, *K*_m AKA_ = 606 μM). Moreover in vitro assays have shown that reconstituted KGDHC complex displays activity towards AKA albeit at much lower catalytic activity and affinity (Nemeria et al., 2018). In order to demonstrate that KGDHC (i.e. OGDH) is the alternative source of glutaryl-CoA in the *GCDH/DHTKD1* double KO, we generated *GCDH/DHTKD1/OGDH* triple KO and *GCDH/OGDH* double KO cell lines. For this, we selected one *GCDH/DHTKD1* double KO and one *GCDH* single KO cell line and targeted *OGDH* using CRISPR-Cas9 (Figures S2G and S2H). OGDH protein was decreased in double and triple KO cell lines, but a variable amount of residual OGDH protein remained detectable, i.e. 9-44% and 36-68% in *GCDH/DHTKD1/OGDH* triple KO and *GCDH/OGDH* double KO cell lines, respectively. This may be attributed to reactivity of the antibody against the α-ketoglutarate dehydrogenase-like protein OGDHL that shares 85% of similarity with OGDH (Bunik et al., 2008) or incomplete KO of *OGDH*. Oxidative decarboxylation of AKG (KGDHC activity) in HEK-293 KO cell lysates was decreased by 55-96% in double and triple KO cell lines proving the successful targeting of *OGDH*. Clones with the lowest activities were selected for further evaluation.

All *GCDH/DHTKD1/OGDH* triple KO cell lines showed a decrease in the level of C5DC to control levels or lower (Figure 2B and Figure S2I). As expected the level of AAA remained increased in all triple KO cells, but with some variation between clones (Figure 2B and Figure S2J). The *GCDH/OGDH* double KO cell lines showed larger variation in the C5DC level, probably due to clonal isolation. Two cell lines had similar C5DC levels as the parental single GCDH KO, one has an intermediate level and one had control levels (Figure 2B and Figure S2I). The level of AAA remained at control level in all cell lines (Figure 2B and Figure S2J) illustrating that the formation of functional KADHC complexes does not depend on OGDH. Combined these cell line data demonstrate that OGDH is responsible for the residual C5DC formation in *GCDH/DHTKD1* double KO cell lines.

### DHTKD1 and OGDH Interact and Share DLST and DLD

We next studied the impact of transient overexpression of the different E1, E2, and E3 subunits on the activity of KGDHC and KADHC. In HEK-293 cell lysates, the oxidative decarboxylation of AKA is approximately 40 times lower than that of AKG (Figure 3A and Fig. S3A). The activity with AKG only increased upon transfection of OGDH, while the activity with AKA increased upon transfection of DHTKD1 and/or OGDH (Figure 3A and Figures S3B and S3C). Similar results were obtained when DLST and DLD were cotransfected with OGDH and DHTKD1. We also assessed enzyme activity after transfection of a novel α-ketoglutarate dehydrogenase-like protein (OGDHL) (Bunik et al., 2008) and MRPS36, a protein that has been implicated in the recruitment of DLD to the KGDHC complex (Heublein et al., 2014). Transient transfection of OGDHL or MRPS36 did not increase ketoacid dehydrogenase activity in HEK-293 cell lysates using either AKA or AKG substrates (Figure S3C). These data further show that DHTKD1 is highly specific for AKA, whereas OGDH has activity with AKA and AKG. Due to this substrate overlap, OGDH is able to compensate for the loss of DHTKD1 function.

**Figure 3.**
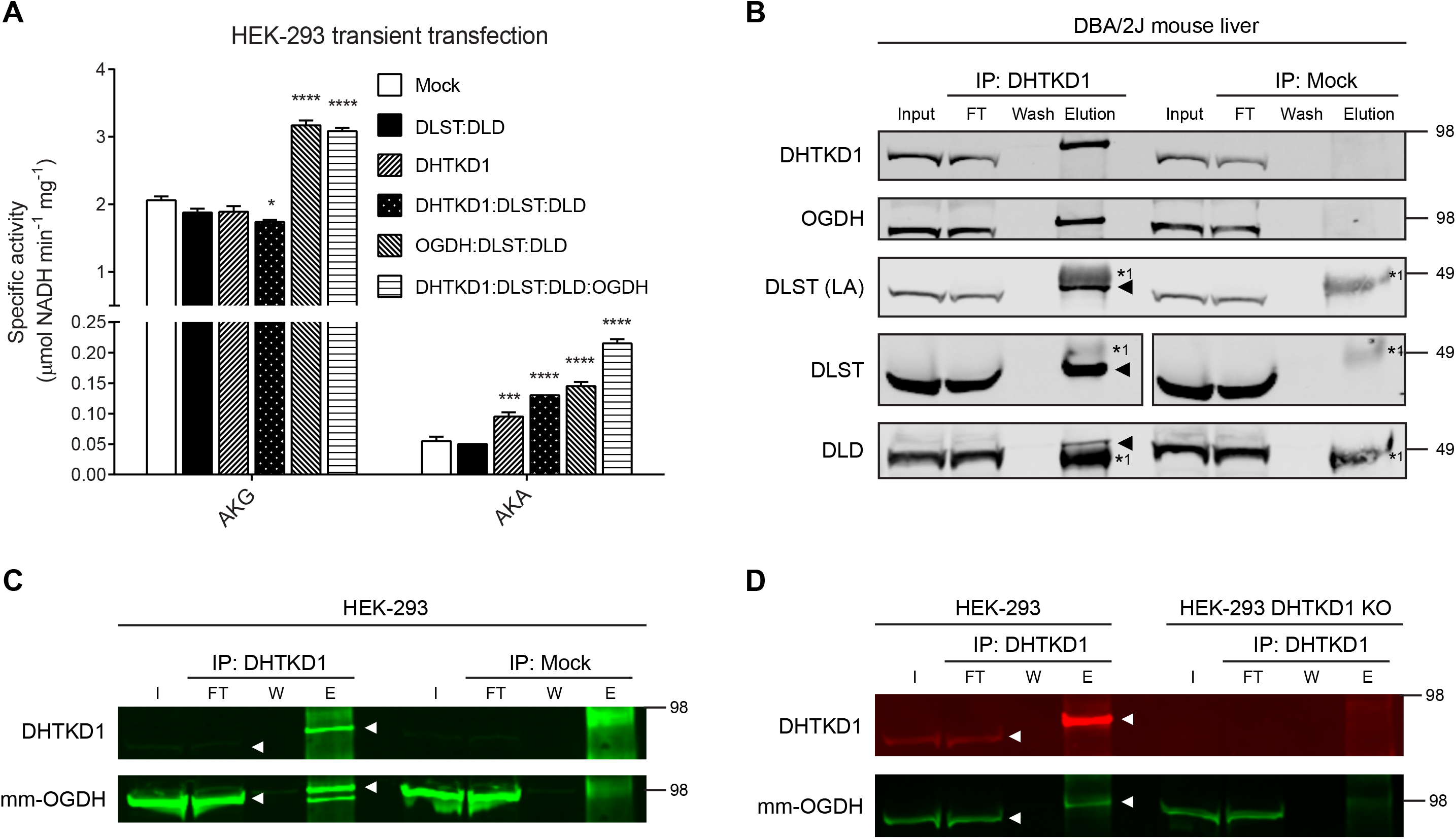
The KADHC Complex Contains DLST, DLD but also OGDH. (A) DHTKD1 and OGDH overexpression leads to increased oxidative decarboxylation of AKA. Activity of AKG and AKA oxidative decarboxylation catalyzed by KGDHC and KADHC complexes in cell lysates of HEK-293 cells upon transient transfection of the indicated genes. HEK-293 cells were transfected with constructs encoding *DHTKD1, DLST, DLD* and *OGDH*. The ratio of DNA between the different gene combinations was maintained constant. *, p<0.05; ***, p<0.001 and ****, p<0.0001. (B) OGDH co-purifies with DHTKD1. DHTKD1 was immunoprecipitated from a liver homogenate of a DBA/2J mouse. DLST (using either an antibody against the lipoic moiety or the protein), DLD and OGDH were confirmed as interacting partners of DHTKD1 in the mammalian KADHC complex. Aliquots of homogenate (Input: 2.5% of total fraction volume), the unbound protein after incubation with IP antibody (FT, flow-through: 2.5% of total fraction volume), the last wash (Wash: 2% of total fraction volume) and eluted fraction (Elution: 100% of total fraction volume) were analyzed by WB by successively using anti-DHTKD1, anti-OGDH, anti-lipoic acid, anti-DLST and anti-DLD antibodies. Heavy (*1) chain of IP antibody is indicated. The triangle indicates the position of the co-IP protein when close to the heavy chain of the IP antibody. The position of molecular mass marker proteins (in kDa) is given. All proteins in the elution fraction migrate slightly higher than the proteins in the lanes containing input and FT likely due to lower protein amount in this lane. (C and D) OGDH is co-immunoprecipitated from HEK-293 cell lysate by DHTKD1 (C) and the anti-DHTKD1 antibody is specific for DHTKD1 and does not immunoprecipitate OGDH (D). DHTKD1 was immunoprecipitated from HEK-293 cell lysate and from a DHTKD1 KO HEK-293 cell lysate. Aliquots of homogenate (I, input: 2.5% of total fraction volume), the unbound protein after incubation with IP antibody (FT, flow-through: 2.5% of total fraction volume), the last wash (W: 2% of total fraction volume) and eluted fraction (E: 100% of total fraction volume) were analyzed by WB by successively using a mouse monoclonal anti-OGDH (mm-OGDH, 66285-1-lg) and anti-DHTKD1. The white triangle indicates the IB band. The position of molecular mass marker proteins (in kDa) is given.

The biochemical properties of the KADHC have not been characterized in mammalian tissues. In order to identify the other members of the complex, we performed immunoprecipitation (IP) experiments. DHTKD1 was successfully immunoprecipitated from DBA/2J mouse liver. Immunoblot analysis showed that DLST and DLD are the interacting E2 and E3 subunits (Figure 3B). Surprisingly, the immunoprecipitate also contained a protein with a molecular weight slightly larger than DHTKD1. Given the molecular weight and the interaction with DLST and DLD, we hypothesized that this could be OGDH. Immunoblotting using OGDH-specific antibodies confirmed this hypothesis (Figure 3B). Additional evidence came from proteomic analysis of gel slices of interest, which identified OGDH, DLST and DLD as interacting partners of DHTKD1 (Figure S3D and Table S1). This result indicates that DHTKD1 and OGDH can form a hybrid KGDHC and KADHC.

Given the homology between DHTKD1 and OGDH, we considered the possibility that the immunoprecipitation of OGDH with the anti-DHTKD1 antibody could be the result of direct binding of the anti-DHTKD1 antibody to OGDH. In an attempt to further confirm the DHTKD1-OGDH interaction, we tried immunoprecipitation of OGDH from DBA/2J mouse liver using two different commercially available OGDH antibodies (Figures S3E-G). A rabbit polyclonal antibody was able to immunoprecipitate OGDH, but we were only able to demonstrate the interaction with DLST, and not with DLD and DHTKD1 (Figures S3E and S3F). An immunoprecipitation of DLST from DBA/2J mouse liver did reveal the formation of stable complexes between DLST, DLD and OGDH and DHTKD1 (Figure S3G), but this result does not necessarily provide additional evidence for the existence of a hybrid complex. Therefore to rule out the potential of unspecific pull-down of OGDH by the anti-DHTKD1 antibody, we performed immunoprecipitation experiments in control HEK-293 cells and *DHTKD1* KO HEK-293 cells. Using the anti-DHTKD1 antibody, OGDH was co-immunoprecipitated in HEK-293 control lysates (Figure 3C), but was not detected in a pull-down from HEK-293 *DHTKD1* KO lysates (Figure 3D). These data demonstrate the specificity of the anti-DHTKD1 antibody and support the existence of a hybrid KGDHC and KADHC containing DHTKD1, OGDH, DLST and DLD.

## DISCUSSION

Mutations in *DHTKD1* lead to a defect in KADHC activity and cause α-aminoadipic and α-ketoadipic aciduria (Danhauser et al., 2012; Hagen et al., 2015), a condition that is currently regarded as a biochemical phenotype of questionable clinical significance, which means that it can be diagnosed through biochemical and genetic methods, but is considered not harmful (Fischer and Brown, 1980; Fischer et al., 1974; Goodman and Duran, 2014). This observation led to the hypothesis that GA1 can be treated through substrate reduction by inhibiting DHTKD1, which was speculated to divert the accumulation of neurotoxic glutaryl-CoA into less harmful accumulation of α-aminoadipic and α-ketoadipic acid. Unexpectedly, *Gcdh/Dhtkd1* double KO mice revealed similar biochemical and clinical outcomes as observed for the existing GA1 mouse model (Biagosch et al., 2017). We confirmed these results by making use of the naturally occurring hypomorphic *Dhtkd1* allele in C57BL/6 and the wild type *Dhtkd1* allele in DBA/2J. These animal studies show that DHTKD1 deficiency does not limit the accumulation of plasma C5DC and urine 3-hydroxyglutaric and glutaric acid in GA1 mice, and point to the existence of an alternative enzymatic source of glutaryl-CoA.

To further investigate the enzymatic origins of glutaryl-CoA in the lysine degradation pathway, we generated CRISPR-Cas9 gene knockouts in HEK-293 cells. We show that in *DHTKD1/GCDH* double KO cell lines the GA1 biomarker C5DC is reduced 2-fold compared with *GCDH* single KO cell lines. The decrease in C5DC suggests that there may be some therapeutic potential for inhibition of DHTKD1, but it also further establishes the existence of a significant alternative source of glutaryl-CoA, as C5DC is not completely reduced to control levels (Figure 4). Different candidate sources of glutaryl-CoA were put forward, one of which involved the formation of glutaryl-CoA via an unknown enzymatic conversion in the pipecolic acid pathway (Figure S1) (Biagosch et al., 2017). Recent studies, however, argue against an important role for the pipecolic acid pathway in lysine degradation (Crowther et al., 2019; Pena et al., 2017; Posset et al., 2015). OGDH and OGDHL, the two paralogs of DHTKD1, are potential candidate sources of glutaryl-CoA. OGDH is the E1 subunit of the KGDHC, a key enzyme from the TCA cycle that catalyzes the conversion of AKG into succinyl-CoA (Sheu and Blass, 1999; Yeaman, 1989). In fact, porcine heart KGDHC and in vitro reconstituted human KGDHC can use AKA as substrate (Nemeria et al., 2017; Nemeria et al., 2018; Sauer et al., 2011), albeit with lower affinity and catalytic efficiency. The KGDHC contains multiple copies of OGDH, DLST and DLD, as the E1, E2 and E3 subunits, respectively (Reed and Hackert, 1990). OGDHL shares high homology with OGDH (Bunik et al., 2008) and is highly expressed in brain, kidney and liver, but its function remains unknown. We now show that overexpression of the individual DHTKD1 and OGDH subunits in HEK-293 cells resulted in increased oxidative decarboxylation of AKA, which argues in favor of OGDH as the source of C5DC in *GCDH/DHTKD1* double KO cells. Indeed knocking out *OGDH* in *GCDH/DHTKD1* double KO cells led to a reduction in C5DC to control or even lower levels. These data firmly establish that OGDH is the enzyme responsible for the partial compensation of deficient or inhibited DHTKD1 activity (Figure 4).

**Figure 4.**
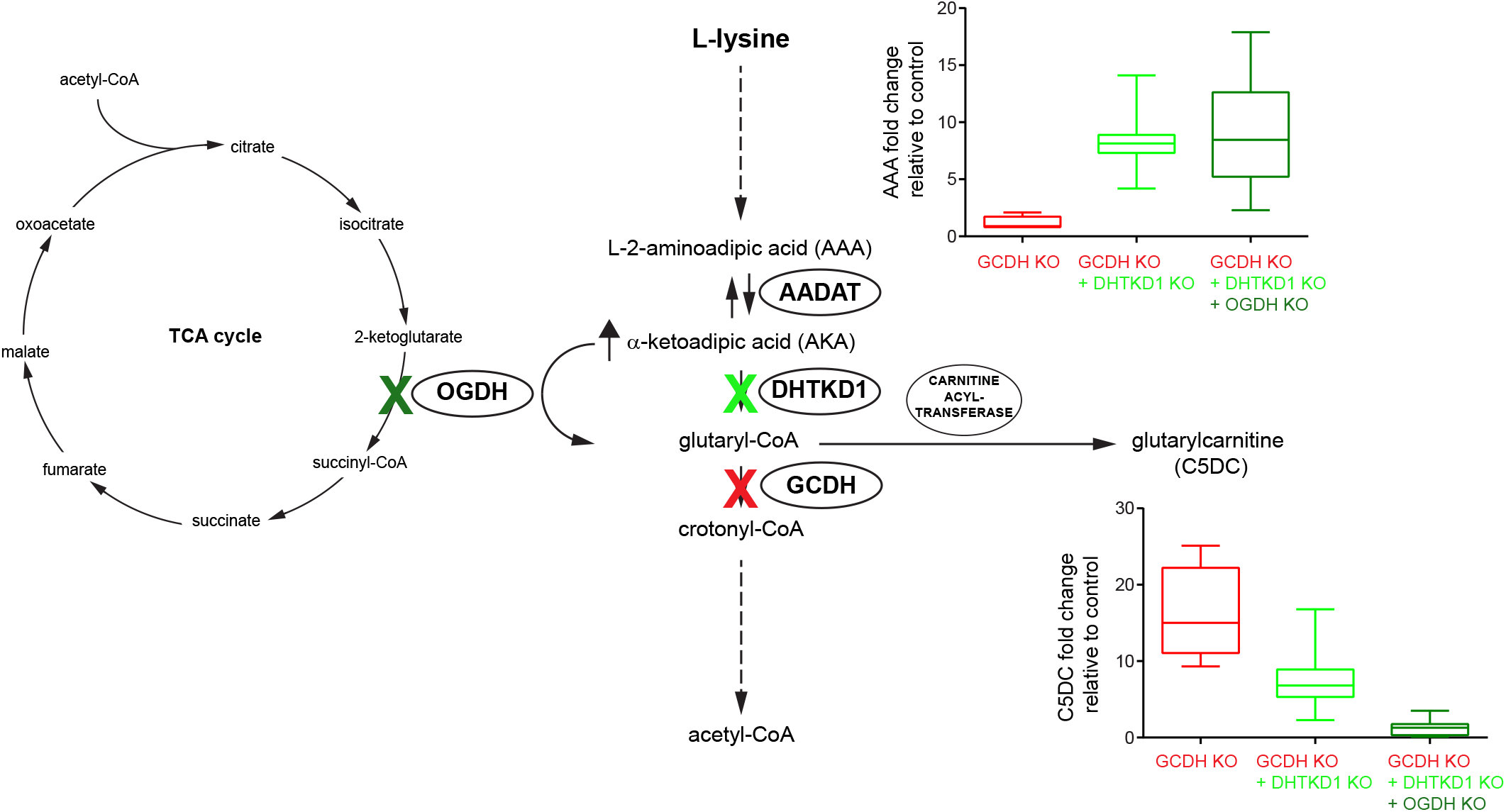
Schematic Representation of the L-Lysine Degradation Pathway upon Inhibition of DHTKD1. Knockout of *GCDH* in HEK-293 cells leads to increased levels of glutaryl-CoA and its metabolite C5DC (in red). This condition mimics the biochemical state occurring in GA1. A substrate reduction therapy for treatment of GA1 by inhibition of DHTKD1 was evaluated by knocking out *DHTKD1* in *GCDH* KO cells (in light green). Knocking out *DHTKD1* increased AAA levels and decreased C5DC levels by approximately 2-fold. Knockout of *OGDH* in *GCDH/DHTKD1* double KO cells further decreased C5DC to wild-type levels (in dark green). High concentrations of AKA such as those that occur after DHTKD1 inhibition or knockout stimulate OGDH activity leading to near normal pathway flux.

We also investigated the composition of the KADHC complex using co-immunoprecipitation of DHTKD1 from mouse liver and kidney, and human HEK-293 cells. We show that DHTKD1 interacts with DLST and DLD, which corroborates the previously reported in vitro assembly of a functional KADHC with DHTKD1, DLST and DLD (Nemeria et al., 2018). Interestingly, we show that DHTKD1 also interacts with OGDH indicating that OGDH and DHTKD1 not only exist as isolated KGDHC and KADHC, but also as a hybrid complex.

As predicted based on the human and mouse genetics data, knocking out *DHTKD1* increased cellular AAA levels. In contrast, knocking out *OGDH* did not change cellular AAA levels confirming that DHTKD1 is relatively specific for AKA, whereas OGDH is more promiscuous and can accept not only AKG, but also AKA. Although OGDH is active with AKA, DHTKD1 deficiency leads to increased levels of AAA and AKA, which indicates that higher substrate levels are necessary to maintain adequate pathway flux. High levels of pathway intermediates may be disadvantageous as it causes increased urinary loss of these metabolites, and can limit overall flux by inhibiting upstream enzymes in the pathway.

We furthermore show that AAA levels are higher in *GCDH/DHTKD1* double KO cells when compared to *DHTKD1* single KO cells (Figure S2F). A similar phenomenon was observed in the *Gcdh* KO mouse. *Gcdh* KO mice had an increase in plasma AAA and urine AKA, which appeared independent of the *Dhtkd1* genotype (Figure 1). This increase in AAA and AKA can be attributed to product inhibition of KGDHC and KADHC by glutaryl-CoA (Nemeria et al., 2018), which is mediated via the shared E2 subunit (DLST) (Sauer et al., 2005). Interestingly, affinity chromatography, co-precipitation and protein complementation assays have shown that GCDH interacts with DLST (Schmiesing et al., 2014). DLST transfers the acyl-TPP intermediate from E1 to its lipoyl group by reductive acylation, followed by transfer of the acyl group to CoA, generating an acyl-CoA (e.g. generation of glutaryl-CoA from 4-carboxy-1-hydroxybutyl-TPP). Close proximity of DLST and GCDH allows efficient channeling of glutaryl-CoA to GCDH for oxidative decarboxylation into crotonyl-CoA and as such limit the product inhibition of DLST. The presence of DHTKD1 and OGDH, two enzymes with overlapping and specific substrate specificities, in a single complex with the shared E2 and E3 subunits, and possibly GCDH increases the complexity of the enzyme complex, but could offer additional possibilities to coordinate metabolic flux in the TCA cycle and the lysine degradation pathway.

In conclusion, our study has identified both physical and functional interactions between DHTKD1 and OGDH. We show that OGDH and DHTKD1 can exist as a novel hybrid α-ketoglutaric and α-ketoadipic acid dehydrogenase complex and display substrate overlap in vivo. The loss of DHTKD1 in GCDH-deficient cells does not prevent the accumulation of GCDH substrates due to the activity of OGDH driven by the elevated levels of AKA. This close functional relationship between DHTKD1 and OGDH limits the therapeutic potential of DHTKD1 inhibition for the treatment of GA1. It is, however, conceivable that DHTKD1 inhibition can limit the production of GCDH substrates under conditions of increased lysine degradation such as catabolic episodes. Alternatively, given the homology between OGDH and DHTKD1 it may be possible to develop a DHTKD1 inhibitor that is also able to partially inhibit OGDH. Both options should be addressed in future studies.

## SUPPLEMENTAL INFORMATION

The Supplemental Information includes three figures and one table and can be found with this article online.

## ACKNOWLEDGMENTS

We thank Dr. Jonna Westover, University of Utah, Dr. Frank Frerman, University of Colorado and Dr. Michael Woontner from the Children’s Hospital Colorado for providing the anti-GCDH antibody, and Dr. Matthew Hirschey for providing the *Gcdh* KO mice. We thank Ethellyn Panta, Hongjie Chen, Jianqiang Zeng, Purvika Patel and Xing Ni for excellent technical assistance with the acylcarnitine and organic acid analyses, and Amelia Gabler for excellent technical assistance with the proteomic analysis. Research reported in this publication was supported by the Eunice Kennedy Shriver National Institute of Child Health & Human Development of the National Institutes of Health under Award Number R03HD092878 (to S.H.) and R21HD088775 (to S.M.H. and to R.J.D.).

## AUTHOR CONTRIBUTIONS

Conceptualization, SMH, RJD; Methodology, JA, RCH, RS, CY; Investigation, JL, TD; Writing – Original Draft, JL; Writing –Review & Editing, SH; Funding Acquisition, SH, RJD.

## STAR★ METHODS

### CONTACT FOR REAGENT AND RESOURCE SHARING

Further information and requests for resources and reagents should be directed to and will be fulfilled by the Lead Contact, Sander M. Houten.

### EXPERIMENTAL MODEL AND SUBJECT DETAILS

#### Mouse Strains

B6.129S4-*Gcdh^tm1Dmk^*/Mmnc mice (Koeller et al., 2002) were provided by Dr. Matthew Hirschey (Duke Molecular Physiology Institute, Duke University Medical Center, Durham, NC) after approval of the Mutant Mouse Resource & Research Centers. All animal experiments were approved by the IACUC of the Icahn School of Medicine at Mount Sinai (#2016-0490) and comply with the National Institutes of Health guide for the care and use of Laboratory animals (NIH Publications No. 8023, revised 1978).

#### Cell Lines

HEK-293 cells (ATCC, Cat#CRL-1573; RRID: CVCL_0045) were cultured in DMEM with 4.5 g/L glucose, 584 mg/L L-glutamine and 110 mg/L sodium pyruvate, supplemented with 10% fetal bovine serum (FBS), 100 U/mL penicillin, 100 μg/mL streptomycin, in a humidified atmosphere of 5% CO_2_ at 37°C.

### METHOD DETAILS

#### Animal Experiments

B6.129S4-*Gcdh^tm1Dmk^*/Mmnc mice (Koeller et al., 2002) were fully backcrossed to C57BL/6NJ (homozygous for WT *Nnt*). *Gcdh*^+/−^ mice were crossed with *Dhtkd1*^D2/B6^ (2 generations C57BL/6N) mice and double heterozygous mice from the resulting F1 were then intercrossed to generate an experimental cohort of B6D2F2 mice. Urine of individual mice was collected on multiple days and pooled in order to obtain sufficient sample volume. Mice were euthanized in a random fed state (afternoon) at 10 weeks of age using CO_2_. Blood from the inferior vena cava was collected for EDTA plasma isolation, and organs were rapidly excised and snap frozen in liquid nitrogen and subsequently stored at −80°C.

#### Generation of CRISPR-Cas9 Knockout Cell Lines

The generation of gene knockout cell lines using the CRISPR-Cas9 genome editing technique was performed essentially as described (Ran et al., 2013), with minor modifications (Violante et al., 2019). For each gene, at least two different guides were chosen and cloned into the pSpCas9(BB)-2A-GFP vector. Following plasmid purification, HEK-293 cells were transfected using lipofectamine 2000. Forty-eight hours after transfection, cells with GFP signal were sorted as single cells into 96 well plates by FACS analysis. The clonal cells were cultured for approximately two weeks and subsequently collected for DNA sequencing and immunoblot analysis (Violante et al., 2019).

We aimed to select multiple independent clonal KO cell lines. HEK-293 cells are near triploid with 62–70 chromosomes/cell making mutation analysis through Sanger sequencing challenging (Bylund et al., 2004; Lin et al., 2014). In order to make the mutation analysis simple, we tried to select clones with apparent homozygous or compound heterozygous mutations (deletions, insertions and indels) encoding a nonsense allele. This implies that cell lines with an apparent homozygous mutation harbored identical triallelic mutations. In compound heterozygous cell lines two alleles must be identical. Mutation detection was performed by direct sequencing of the genomic DNA. Primers were chosen to flank the region surrounding the respective guides for each gene (PCR products between 250 and 500 bp). Primer sequences are available upon request. We were not able to resolve the exact mutation for all used clonal cell lines, but all mutations were demonstrated to be deleterious through immunoblotting and functional assays.

#### Immunoblotting

Cells were lysed in RIPA buffer supplemented with protease inhibitors cocktail (Pierce), centrifuged 10 min at 12,000 ×*g*, 4°C and total protein determined by the BCA method. Proteins were separated on a Bolt™ 4–12% Bis-Tris Plus Gel, blotted onto a nitrocellulose membrane and detected using the following primary antibodies: DHTKD1 (GTX32561, Genetex), OGDH (66285-1-Ig, Proteintech), GCDH (gift of Michael Woontner, Children’s Hospital Colorado) and citrate synthase [GT1761] (GTX628143, Genetex) or [N2C3] (GTX110624; RRID:AB_1950045, Genetex). Proteins were visualized using IRDye 800CW or IRDye 680RD secondary antibodies (LI-COR, 926-32210; RRID:AB_621842, 926-68070; RRID:AB_10956588, 926-32211; RRID:AB_621843, and 926-68071; RRID:AB_10956166) and Image Studio Lite software (version 5.2, LI-COR). Equal loading was checked by Ponceau S staining and the citrate synthase signal.

#### Metabolite Analyses

Plasma acylcarnitines, plasma amino acids and urine organic acids were measured by the Mount Sinai Biochemical Genetic Testing Lab (now Sema4). Urine organic acids were quantified using a standard curve and pentadecanoic acid as internal standard.

For quantification of α-aminoadipic acid (AAA) and glutarylcarnitine (C5DC) in HEK-293 KO, the cell lines were incubated in supplemented DMEM with 0.4 mM L-carnitine for 24 h. Cells were collected, washed in PBS, flash-frozen in liquid nitrogen and stored at −80°C until further use. Samples were sonicated in ice cold demineralized water and then deproteinized by adding ice cold acetonitrile up to 80% (v/v). After 10 min centrifugation at maximum speed (4°C), the supernatant was dried under nitrogen and the residue was dissolved in 75 μL of demineralized water. Acylcarnitine internal standards were obtained from Cambridge Isotope Laboratories (Andover, MA, USA) (NSK-B-1, containing ^2^H_9_-carnitine (d9-C0), ^2^H_3_-acetylcarnitine (d3-C2), ^2^H_3_-propionylcarnitine (d3-C3), ^2^H_3_-butyrylcarnitine (d3-C4), ^2^H_9_-isovalerylcarnitine (d9-C5), ^2^H_3_-octanoylcarnitine (d3-C8), ^2^H_9_-myristoylcarnitine (d9-C14) and ^2^H_3_-palmitoylcarnitine (d3-C16) and dissolved in 40 mL LC-MS-grade methanol. QCs and samples were prepared by mixing 20 μL of each with 100 μL of internal standard solution. Samples were centrifuged at 10,000 rpm, 10 min. 20 μL of the supernatant was transferred to a 96-well microplate and evaporated under a stream of nitrogen. Butylation was performed by adding 50 μl of 3N n-butanol-HCl at 65°C for 15 min. After drying under nitrogen, the samples were reconstituted in 200 μl of methanol:water (8:2 v/v) and injected into the LC-MS/MS system. Amino acids were measured by LC-MS/MS as described (Le et al., 2014). Protein pellets were dissolved in 25 mM KOH, 0.1% Triton X-100, and the protein amount determined by the BCA method for subsequent normalization of the amino acid and acylcarnitines levels.

#### Immunoprecipitation

Frozen mouse tissues (liver and kidney) were homogenized in ice-cold lysis buffer (50 mM MOPS, pH 7.4, 0.1% Triton X-100: ~20 mg per 1 mL lysis buffer) using a TissueLyser II apparatus (Qiagen). HEK-293 cell pellets were sonicated in ice-cold lysis buffer. Protein A or G magnetic Surebeads (Bio-Rad) were used for co-immunoprecipitations depending on the affinity for the specific antibody. Antibodies were incubated with appropriate magnetic beads for 10 min at RT and beads washed to remove unbound antibody. Lysates were centrifuged at 1,000 ×*g*, 10 min at 4°C, and pre-cleared with an unrelated antibody of the same isotype as the antibody used for IP. Pre-cleared lysates were incubated with 5 μg of the specific IP antibody bound beads or a mock control antibody for 2 h at 4°C under gentle rotation. Immunoprecipitates were washed in PBS + 0.1% Tween 20, resuspended in 2× SDS-PAGE loading buffer (Bolt LDS sample buffer and sample reducing agent), boiled for 10 min at 70°C and analyzed by western blot. Anti-DHTKD1 (GTX32561, Genetex), anti-lipoic acid (437695; RRID:AB_212120, Calbiochem), anti-GCDH (gift of Michael Woontner, Children’s Hospital Colorado), anti-DLD (G-2) (sc-365977; RRID:AB_10917587, Santa Cruz), anti-OGDH (GTX33374, Genetex), anti-OGDH (66285-1-lg, Proteintech), anti-DLST (ab110306; RRID:AB_10862702, Abcam) and mock anti-ABCD3 (TA310030, Origene) or anti-DCPS (ab57314; RRID:AB_941255, Abcam) antibodies were used for IP or western blot as indicated. Total protein staining was obtained using the REVERT total protein stain kit (LI-COR) and Quick Western Kit - IRDye^®^ 680RD (LI-COR) was used to avoid detection of the IP antibody, when appropriate.

#### Mass Spectrometry Analysis

Forty μL of co-IP eluted samples were loaded on 4–12% Bolt gel (Invitrogen), stained with Coomassie SimplyBlue SafeStain (ThermoFisher) and submitted to the Microchemistry and Proteomics Core Facility at Memorial Sloan Kettering Cancer Center for in-gel tryptic digestion and protein identification by mass spectrometry. Scaffold 4.8.4 (Proteome Software Inc., Portland, Oregon) was used to validate MS/MS-based peptide and protein identifications. Peptides were identified from MS/MS spectra by searching against a library of known/likely interactors of DHTKD1.

#### Overexpression in HEK-293 Cells

Human cDNA clone for *DHTKD1* (NM_018706) was obtained from Origene (RC200967), encoding a C-terminal tagged DHTKD1-Myc-DDK on a pCMV6 plasmid. One rare variant (L308) was detected on *DHTKD1* cDNA and was therefore replaced by the common variant R308 by site directed mutagenesis using the following forward and reverse primers: 5’-GGC AGC AGT CTC **G**CC AAG ACG GCG-3’ and 5’-CGC CGT CTT GG**C** GAG ACT GCT GCC-3’ (mutagenic nucleotides in bold). In order to obtain the untagged protein, a STOP codon was introduced before the C-terminal tags by site directed mutagenesis using as template pCMV6-DHTKD1(R308)-Myc-DDK and the following forward and reverse primers: 5’-CCA AGA CCT TCG CT**T AA**C GTA CGC GGC CGC-3’ and 5’-GCG GCC GCG TAC G**TT A**AG CGA AGG TCT TGG-3’ (mutagenic nucleotides in bold). However, no impact on activity was detected for the presence of the variant or the C-terminal tag (Fig. S3A).

Human cDNA clones for *OGDH* (BC004964), *DLST* (BC001922), *DLD* (BC018696) and *MRPS36 (KGD4*) (BC015966) were provided by TransOMIC technologies (Huntsville, AL). The genes were amplified by using High Fidelity PrimeSTAR GXL Polymerase (Takara) and cloned into AsiSI(SgfI)/MluI cloning site of the pCMV6-Entry vector (Origene). Each construct was analyzed by restriction digestion and sequencing. Human cDNA clone for *OGDHL* (NM_018245) was obtained from Origene (RC205225), encoding a C-terminal tagged OGDHL-Myc-DDK on a pCMV6 plasmid. Two rare variants (L511 and N573) were detected on *OGDHL* cDNA and were therefore replaced by the common variant P511 and D573 by site directed mutagenesis using the following forward and reverse primers: 5’-CCC ATG TTC ACC CAG C**C**G CTC ATG TAC AAG CAG-3’, 5’-CTG CTT GTA CAT GAG C**G**G CTG GGT GAA CAT GGG-3’ and 5’-ATA AAG CAC TGG TTG **G**AC TCC CCC TGG CCT G-3’, 5’-CAG GCC AGG GGG AGT **C**CA ACC AGT GCT TTA T-3’, respectively (mutagenic nucleotides in bold). Cell transfection grade plasmid DNA was obtained for every plasmid construct with NucleoBond Xtra Midi Plus EF DNA purification kit (Macherey-Nagel).

HEK-293 cells were transfected with different combinations of vectors performing a total of 6.9 μg/T-25 flask using lipofectamine 2000 as transfection agent and Opti-MEM I Reduced Serum (Gibco). After 24 h, cells were harvested, washed and flash-frozen in liquid nitrogen. Pellets were stored at −80°C until further use for immunoblot and/or enzyme activity analysis.

#### KGDHC and KADHC Activity

HEK-293 cells were lysed in 50 mM MOPS, pH 7.4, 0.1% Triton X-100, sonicated and centrifuged for 10 min at 1,000 ×*g*, 4°C. The supernatant (100 μg total protein) was mixed with assay buffer (final concentration: 50 mM MOPS, pH 7.4, 0.2 mM MgCl_2_, 0.01 mM CaCl_2_, 0.3 mM TPP, 0.12 mM Coenzyme-A, 2 mM NAD, 2.6 mM β-mercaptoethanol). The reaction was started by the addition of 1 mM substrates: AKG and AKA. The activity of the OGDH and DHTKD1 complexes was followed by measuring the NADH production at 340 nm at 30°C and steady-state velocities were taken from the linear portion of the time curve.

The activity of KGDHC from porcine heart (Sigma K1502) was measured as described above using 1 mM AKG or AKA as substrate.

#### Quantification and Statistical Analysis

Data are displayed as the mean ± standard deviation (SD). One-way analysis of variance with Bonferroni’s multiple comparison test and two-way analysis of variance as used for statistical analysis using GraphPad Prism 7 software (GraphPad, San Diego). Significance is indicated as follows: *, p < 0.05; **, p < 0.01; ***, p < 0.001 and ****, p < 0.0001.

