## Supplementary material for "DHTKD1 and OGDH display in vivo substrate overlap and form a hybrid ketoacid dehydrogenase complex": Figure S1

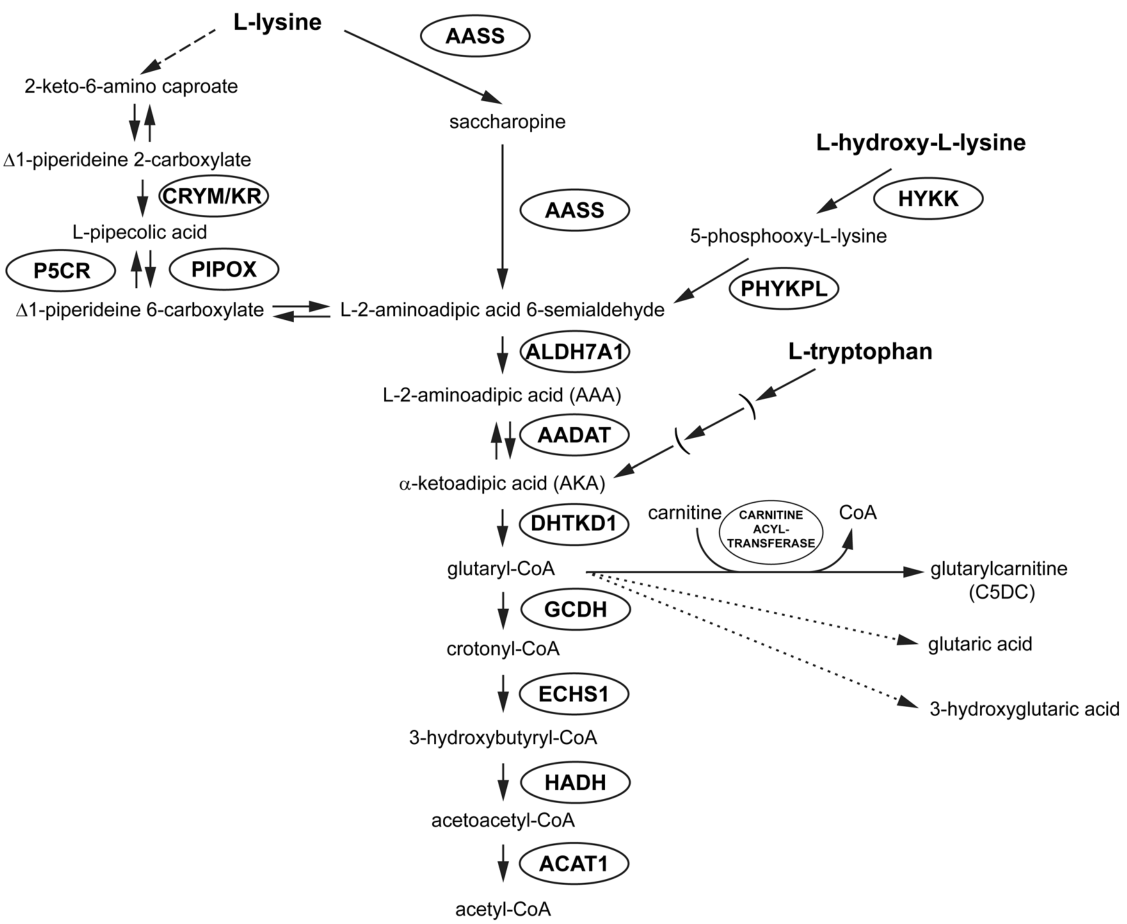


**Figure S1. The lysine degradation pathway. (Related to Fig. 1)**
The degradation of L-lysine may occur via two pathways: the saccharopine and the pipecolic acid pathway. The mitochondrial saccharopine pathway is well established and is the major pathway. The pipecolic acid pathway, which is thought to be more active in the brain, proceeds via oxidative deamination with L-pipecolic acid as intermediate. Degradation of L-hydroxy-L-lysine leads to L-2-aminoadipic acid 6-semialdehyde and cytosolic degradation of tryptophan with kynurenine as intermediate leads to the production of AKA. The intermediate AKA is metabolized into glutaryl-CoA via oxidative decarboxylation by the DHTKD1, the E1 subunit of the α-ketoadipic acid dehydrogenase complex (KADHC). Glutaryl-CoA is converted into crotonyl-CoA via oxidative decarboxylation by GCDH. The GCDH protein is deficient in GA1 leading to the accumulation of glutaryl-CoA metabolites: glutarylcarnitine (C5DC), glutaric acid and 3-hydroxyglutaric acid. The level of AAA correlates with the amount of AKA by reversible transamination catalyzed by AADAT. C5DC is the product of detoxification of accumulating glutaryl-CoA through conjugation with carnitine. AASS, alpha-aminoadipic semialdehyde synthase; ALDH7A1, alpha-aminoadipic semialdehyde dehydrogenase; AADAT, kynurenine/alpha-aminoadipate aminotransferase; CRYM/KR, μ-crystallin/ketimine reductase; DHTKD1, dehydrogenase E1 and transketolase domain containing 1; GCDH, glutaryl-CoA dehydrogenase; ECHS1, enoyl-CoA hydratase; HADH, hydroxyacyl-coenzyme A dehydrogenase; ACAT1, acetyl-CoA acetyltransferase; P5CR, pyrroline-5-carboxylate reductase; PIPOX, peroxisomal sarcosine oxidase; HYKK, hydroxylysine kinase; PHYKPL, 5-phosphohydroxy-L-lysine phospho-lyase. Dashed arrows represent not fully characterized enzymatic steps.
