## Supplementary material for "DHTKD1 and OGDH display in vivo substrate overlap and form a hybrid ketoacid dehydrogenase complex": Figure S2

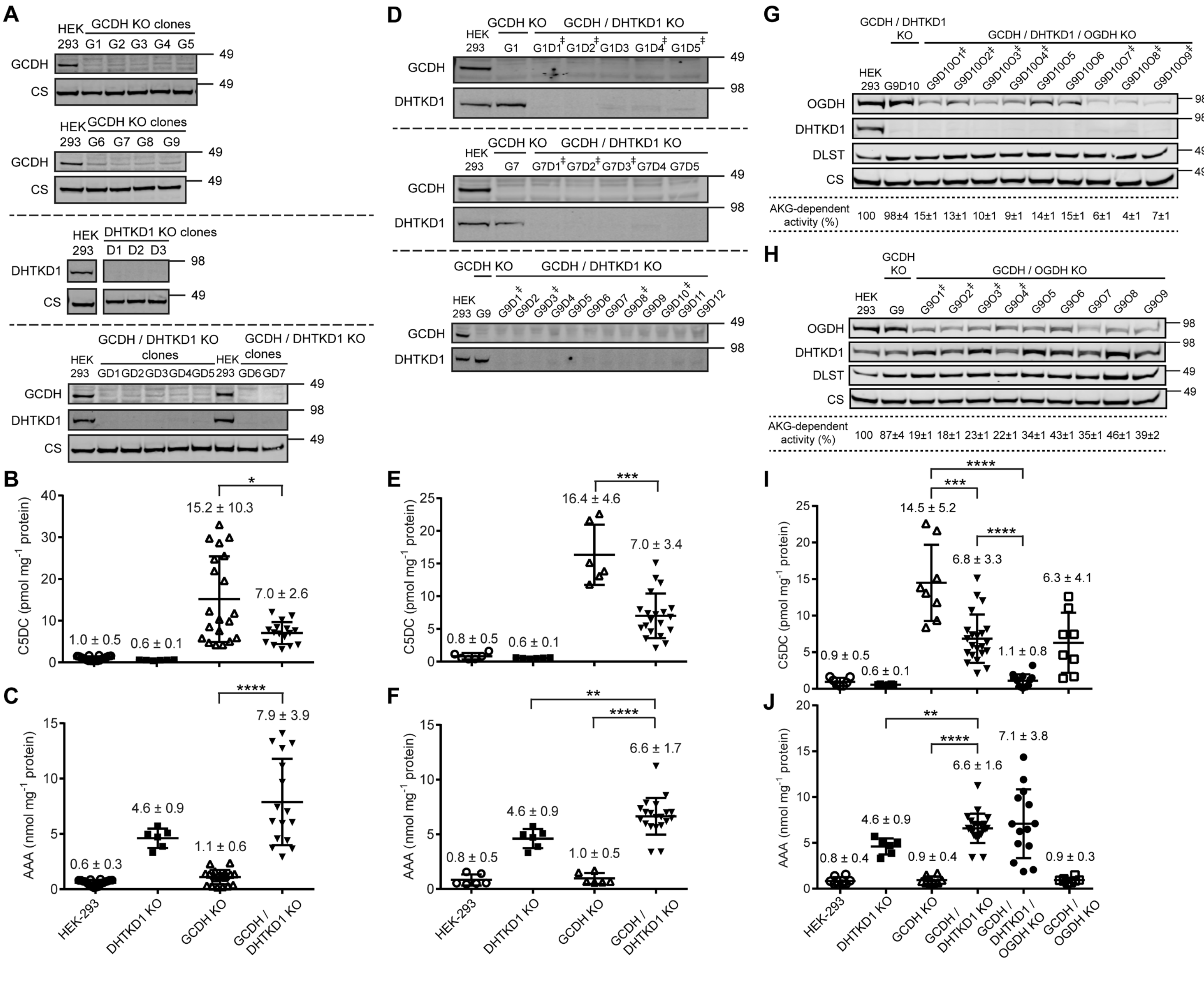
A
HEK 293
GCDH KO clones
G1 G2 G3 G4 G5
G6 G7 G8 G9
GCDH
CS
DHTKD1 KO clones
D1 D2 D3
DHTKD1
GCDH / DHTKD1 KO clones
GD1 GD2 GD3 GD4 GD5 GD6 GD7
49
98
B
C5DC (pmol mg-1 protein)
1.0 ± 0.5
0.6 ± 0.1
15.2 ± 10.3
7.0 ± 2.6
C
AAA (nmol mg-1 protein)
0.6 ± 0.3
4.6 ± 0.9
1.1 ± 0.6
7.9 ± 3.9
HEK-293
DHTKD1 KO
GCDH KO
GCDH / DHTKD1 KO
D
GCDH KO
GCDH / DHTKD1 KO
G1 G1D1‡ G1D2‡ G1D3 G1D4‡ G1D5‡
G7 G7D1‡ G7D2‡ G7D3‡ G7D4 G7D5
G9 G9D1‡ G9D2 G9D3‡ G9D4 G9D5 G9D6 G9D7 G9D8‡ G9D9 G9D10‡ G9D11 G9D12
E
0.8 ± 0.5
0.6 ± 0.1
16.4 ± 4.6
7.0 ± 3.4
F
0.8 ± 0.5
4.6 ± 0.9
1.0 ± 0.5
6.6 ± 1.7
G
GCDH / DHTKD1 KO
GCDH / DHTKD1 / OGDH KO
G9D10 G9D10O1‡ G9D10O2‡ G9D10O3‡ G9D10O4‡ G9D10O5 G9D10O6 G9D10O7‡ G9D10O8‡ G9D10O9‡
OGDH
DHTKD1
DLST
CS
AKG-dependent activity (%)
100 98±4 15±1 13±1 10±1 9±1 14±1 15±1 6±1 4±1 7±1
H
GCDH KO
GCDH / OGDH KO
G9 G9O1‡ G9O2‡ G9O3‡ G9O4‡ G9O5 G9O6 G9O7 G9O8 G9O9
AKG-dependent activity (%)
100 87±4 19±1 18±1 23±1 22±1 34±1 43±1 35±1 46±1 39±2
I
0.9 ± 0.5
0.6 ± 0.1
14.5 ± 5.2
6.8 ± 3.3
1.1 ± 0.8
6.3 ± 4.1
J
0.8 ± 0.4
4.6 ± 0.9
0.9 ± 0.4
6.6 ± 1.6
7.1 ± 3.8
0.9 ± 0.3
GCDH / DHTKD1 / OGDH KO
GCDH / OGDH KO

**Figure S2. *DHTKD1* KO reduces glutarylcarnitine in simultaneously and sequentially generated *GCDH/DHTKD1* double KO HEK-293 cell lines whereas OGDH KO reduces glutarylcarnitine to basal levels in HEK-293 GCDH/DHTKD1/OGDH triple KO cell lines. (Related to Fig. 2)**

(A) Immunoblot verification of *GCDH* single KO (*upper panel*), *DHTKD1* single KO (*middle panel*) and simultaneously generated *GCDH/DHTKD1* double KO (*lower panel*) clonal cell lines. Single *GCDH* KO clones are indicated as G#, single *DHTKD1* KO as D# and simultaneous double *GCDH/DHTKD1* KO clones as GD#. Citrate synthase (CS) was used as loading control. The position of molecular mass marker proteins (in kDa) is given.

(B and C) Biochemical analyses of C5DC (B) and AAA (C) levels in HEK-293 *GCDH/DHTKD1* simultaneous double KO (▼) compared with HEK-293 *DHTKD1* single KO (■), HEK-293 *GCDH* single KO (△) and wild-type HEK-293 cells (○).

(D) Immunoblot verification of *GCDH/DHTKD1* double KOs generated from G1 (*upper panel*), G7 (*middle panel*) and G9 (*lower panel*). Single *GCDH* KO clones are indicated as G# and consecutively generated double *GCDH/DHTKD1* knockout clones as G#D#. The position of molecular mass marker proteins (in kDa) is given. ‡ indicates selected clones to measure the levels of C5DC and AAA.

(E and F) Global statistical analyses of C5DC (E) and AAA (F) levels in HEK-293 *GCDH/DHTKD1* double KO (▼) compared with HEK-293 *DHTKD1* single KO (■), HEK-293 *GCDH* single KO (△) and wild-type HEK-293 cells (○).

(G and H) Immunoblot verification of triple *GCDH/DHTKD1/OGDH* KO clones generated from G9D10 (G) and of double *GCDH/OGDH* KO clones generate from G9 (H). Single *GCDH* KO clones are indicated as G#, double *GCDH/DHTKD1* KO clones as G#D#, double *GCDH/OGDH* KO clones as G#O# and triple *GCDH/DHTKD1/OGDH* KO clones as G#D#O#. The position of molecular mass marker proteins (in kDa) is given. ‡ indicates selected clones to measure the levels of C5DC and AAA. The oxidative decarboxylation of AKG in cell lysates of HEK-293 KOs was measured to evaluate *OGDH* knockout efficiency and is represented as a percentage relative to wild-type activity in HEK-293 cell lysates.

(I and J) Global statistical analyses of C5DC (I) and AAA (J) levels in HEK-293 *GCDH*/*DHTKD1*/*OGDH* triple KO (●) and HEK-293 *GCDH*/*OGDH* double KO (□) compared with HEK-293 *GCDH*/*DHTKD1* double KO (▼), HEK-293 *GCDH* single KO (△), HEK-293 *DHTKD1* single KO (■) and wild-type HEK-293 cells (○).

Error bars indicate SD and the mean ± SD of C5DC and AAA levels are indicated above each group. *, $p<0.05$, **, $p<0.01$; ***, $p<0.001$ and ****, $p<0.0001$.
