## Supplementary material for "DHTKD1 and OGDH display in vivo substrate overlap and form a hybrid ketoacid dehydrogenase complex": Figure S3

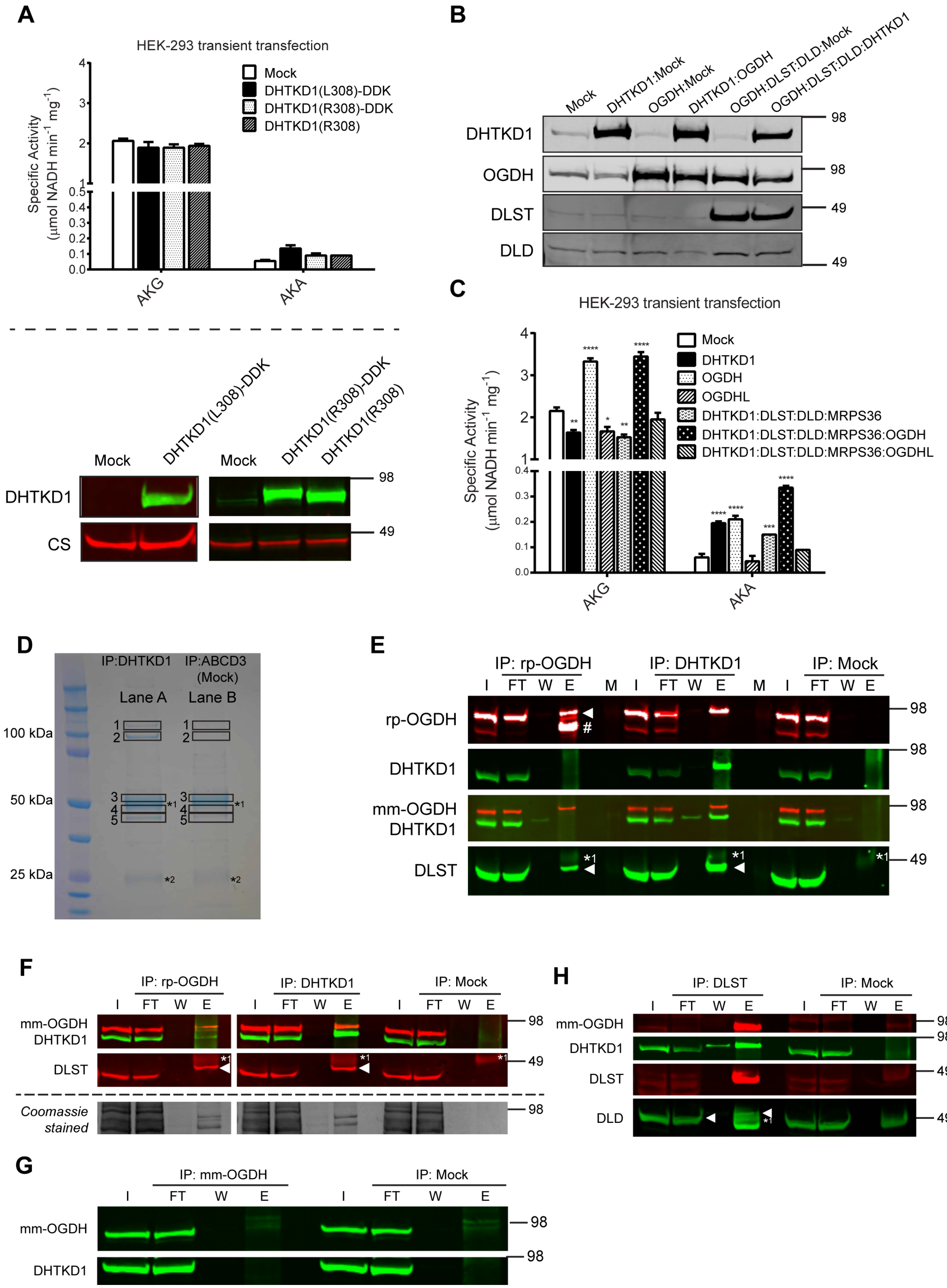
A
HEK-293 transient transfection
Mock
DHTKD1(L308)-DDK
DHTKD1(R308)-DDK
DHTKD1(R308)
Specific Activity
(µmol NADH min-1 mg-1)
AKG
AKA
Mock
DHTKD1(L308)-DDK
Mock
DHTKD1(R308)-DDK
DHTKD1(R308)
DHTKD1
CS
98
49
B
Mock
DHTKD1:Mock
OGDH:Mock
DHTKD1:OGDH
OGDH:DLST:DLD:Mock
OGDH:DLST:DLD:DHTKD1
DHTKD1
OGDH
DLST
DLD
98
98
49
49
C
HEK-293 transient transfection
Mock
DHTKD1
OGDH
OGDHL
DHTKD1:DLST:DLD:MRPS36
DHTKD1:DLST:DLD:MRPS36:OGDH
DHTKD1:DLST:DLD:MRPS36:OGDHL
Specific Activity
(µmol NADH min-1 mg-1)
AKG
AKA
D
IP:DHTKD1
IP:ABCD3
(Mock)
Lane A
Lane B
100 kDa
50 kDa
25 kDa
E
IP: rp-OGDH
IP: DHTKD1
IP: Mock
I FT W E M I FT W E M I FT W E
rp-OGDH
DHTKD1
mm-OGDH
DHTKD1
DLST
98
98
98
49
F
IP: rp-OGDH
IP: DHTKD1
IP: Mock
I FT W E I FT W E I FT W E
mm-OGDH
DHTKD1
DLST
Coomassie
stained
98
49
98
G
IP: mm-OGDH
IP: Mock
I FT W E I FT W E
mm-OGDH
DHTKD1
98
98
H
IP: DLST
IP: Mock
I FT W E I FT W E
mm-OGDH
DHTKD1
DLST
DLD
98
98
49
49

**Figure S3. Overexpression of DHTKD1 and OGDH increase the oxidative decarboxylation of α-ketoadipic acid and immunoprecipitation of DHTKD1, OGDH and DLST from DBA/2J mouse liver or kidney.(related to Related to Fig. 3).**

(A) (*upper panel*) Oxidative decarboxylation of AKG and AKA in cell lysates of HEK-293 cells upon transient transfection with a rare *DHTKD1* variant (L308), the common variant (R308) and without C-terminal tags. 6.9 μg of each plasmid construct was used for transfection. (lower *panel*) Immunoblot of HEK-293 transient transfections depicted on the upper panel. Citrate synthase (CS) was used as loading control and DDK denotes the C-terminal FLAG tag. The position of molecular mass marker proteins (in kDa) is given.

(B) Immunoblot of HEK-293 transient transfections depicted on Fig. 3A and Fig. S3C. The position of molecular mass marker proteins (in kDa) is given.

(C) Oxidative decarboxylation of AKG and AKA in cell lysates of HEK-293 cells upon transient transfection of the indicated genes: *DHTKD1*, *DLST*, *DLD*, *OGDH*, *OGDHL* and *MRPS36*. The amount of DNA has been maintained constant as follows: 6.9 μg in Mock, DHTKD1, OGDH and OGDHL conditions; 1.73 μg of each construct in DHTKD1:DLST:DLD:MRPS36 condition; and 1.38 μg of each construct in DHTKD1:DLST:DLD:MRPS36:OGDH and DHTKD1:DLST:DLD:MRPS36:OGDHL conditions.

(D) Coomassie-stained SDS-PAGE gel of DHTKD1 interacting proteins (lane A) and negative control (lane B). Gel pieces (1 through 5) were cut out for mass spectrometry-based protein identification. *[1] and *[2] indicate the heavy and light chain of the IP antibody, respectively. The position of molecular mass marker proteins (in kDa) is given. Proteins specifically identified in each gel slice are shown in Table S1.

(E) OGDH was immunoprecipitated from liver homogenate of DBA/2J mouse using a rabbit polyclonal anti-OGDH antibody (rp-OGDH, GTX33374). (*upper panel*) The triangle indicates the position of OGDH and # indicates unspecific immunoprecipitation of a protein that was identified as plasminogen by mass spectrometry and is not detected by the monoclonal anti-OGDH antibody. (*lower panel*). The triangle indicates the position of the co-IP protein (DLST) close to the heavy chain of the IP antibody.

(F) OGDH was immunoprecipitated from kidney homogenate of DBA/2J mouse using a rabbit polyclonal anti-OGDH antibody (rp-OGDH, GTX33374). Kidney has a lower expression of plasminogen when compared with liver. Immunoblot analysis showed DLST interacting with OGDH. A band was detected using the anti-DHTKD1 antibody, although with a slightly lower molecular weight than expected for DHTKD1. The triangle indicates the position of the co-IP protein (DLST) close to the heavy chain of the IP antibody.

(G) A third attempt to efficiently immunoprecipitate OGDH from DBA/2J mouse liver homogenate was performed using a monoclonal anti-OGDH antibody (mm-OGDH, 66285-1-Ig). No OGDH was detected in the eluted fraction.

(H) DLST was immunoprecipitated from DBA/2J mouse liver homogenate. The triangle indicates the position of the co-IP protein (DLD) close to the heavy chain of the IP antibody. The position of molecular mass marker proteins (in kDa) is given.

Aliquots of homogenate (I, input: 2.5% of total fraction volume), the unbound protein after incubation with IP antibody (FT, flow-through: 2.5% of total fraction volume), the last wash (W: 2% of total fraction volume) and eluted fraction (E: 100% of total fraction volume) were analyzed by WB by successively using a rabbit polyclonal anti-OGDH (GTX33374) or mouse monoclonal anti-OGDH (mm-OGDH, 66285-1-Ig), anti-DHTKD1, anti-DLST antibodies or Coomassie stained SDS-PAGE gel (E-G) or by successively using anti-DHTKD1, mouse monoclonal anti-OGDH, anti-DLST and anti-DLD antibodies (H). Heavy (*1) chain of IP antibody is indicated.
