## Supplementary material for "DHTKD1 and OGDH display in vivo substrate overlap and form a hybrid ketoacid dehydrogenase complex": Table S1

**Table S1. Mass spectrometric analysis of interacting proteins identified by IP of DHTKD1 from DBA/2J mouse liver. Related to Fig. 3B.**

| Gel slice # | Protein Name | Protein accession number | MW[a] | # unique peptides | | # exclusive spectra | | % sequence coverage | |
|---|---|---|---|---|---|---|---|---|---|
| | | | | Lane A | Lane B | Lane A | Lane B | Lane A | Lane B |
| 1 | 2-oxoglutarate dehydrogenase | Q60597 (ODO1_MOUSE) | 116 kDa (112) | 7 | 0 | 24 | 0 | 7.2 | - |
| | Probable 2-oxoglutarate dehydrogenase E1 component DHKTD1 | A2ATU0 (DHTK1_MOUSE) | 103 kDa (100) | 2 | 0 | 3 | 0 | 2.0 | - |
| 2 | 2-oxoglutarate dehydrogenase | Q60597 (ODO1_MOUSE) | 116 kDa (112) | 3 | 0 | 3 | 0 | 2.5 | - |
| | Probable 2-oxoglutarate dehydrogenase E1 component DHKTD1 | A2ATU0 (DHTK1_MOUSE) | 103 kDa (100) | 17 | 0 | 60 | 0 | 16.6 | - |
| | Dihydrolipoyllysine-residue succinyltransferase component of 2-oxoglutarate dehydrogenase complex | Q9D2G2 (ODO2_MOUSE) | 49 kDa (41) | 2 | 0 | 2 | 0 | 4.6 | - |
| Gel slice # | Protein Name | Protein accession number | MW[a] | # unique peptides | | # exclusive spectra | | % sequence coverage | |
| | | | | Lane A | Lane B | Lane A | Lane B | Lane A | Lane B |
| 3 | Probable 2-oxoglutarate dehydrogenase E1 component DHKTD1 | A2ATU0 (DHTK1_MOUSE) | 103 kDa (100) | 3 | 0 | 3 | 0 | 3.1 | - |
| | Dihydrolipoyl dehydrogenase | O08749 (DLDH_MOUSE) | 54 kDa (50) | 19 | 3 | 57 | 3 | 39.9 | 7.5 |
| | Dihydrolipoyllysine-residue succinyltransferase component of 2-oxoglutarate dehydrogenase complex | Q9D2G2 (ODO2_MOUSE) | 49 kDa (41) | 12 | 4 | 43 | 4 | 32.8 | 8.8 |
| 4 | Probable 2-oxoglutarate dehydrogenase E1 component DHKTD1 | A2ATU0 (DHTK1_MOUSE) | 103 kDa (100) | 3 | 0 | 3 | 0 | 2.9 | - |
| | Dihydrolipoyl dehydrogenase | O08749 (DLDH_MOUSE) | 54 kDa (50) | 4 | 0 | 4 | 0 | 8.3 | - |
| | Dihydrolipoyllysine-residue succinyltransferase component of 2-oxoglutarate dehydrogenase complex | Q9D2G2 (ODO2_MOUSE) | 49 kDa (41) | 8 | 0 | 32 | 0 | 17.8 | - |
| 5 | Probable 2-oxoglutarate dehydrogenase E1 component | A2ATU0 (DHTK1_MOUSE) | 103 kDa (100) | 1 | 0 | 1 | 0 | 1.1 | - |

| | | | | | | | | | |
|---|---|---|---|---|---|---|---|---|---|
| | DHKTD1 | | | | | | | | |
| | Dihydrolipoyl dehydrogenase | O08749 (DLDH_MOUSE) | 54 kDa (50) | 2 | 0 | 2 | 0 | 3.7 | - |
| | Dihydrolipoyllysine-residue succinyltransferase component of 2-oxoglutarate dehydrogenase complex | Q9D2G2 (ODO2_MOUSE) | 49 kDa (41) | 2 | 0 | 3 | 0 | 4.4 | - |
| | Glutaryl-CoA dehydrogenase | Q60759 (GCDH_MOUSE) | 49 kDa (44) | 1 | 2 | 1 | 2 | 3.2 | 5.9 |

Results represent the proteins specifically identified in each gel slice when peptides were run against a library of proteins likely to interact with DHTKD1 (Protein accession numbers: A2ATU0, Q60597, E9Q7L0, Q9D2G2, O08749, Q60759, Q8BHG1, Q9CQX8 and Q9DCX2. Peptides were selected using a stringent filter (99.0% protein threshold, 1 min. peptide and 95% peptide threshold) to avoid false positives. Sequence coverage represents the percentage of sequence (full length protein) matching with peptides found in the analysis. [a]Molecular weight of processed form of the protein (without mitochondrial transit peptide) is shown in parenthesis.
